# Consistent bacterial responses to land use change across the tropics

**DOI:** 10.1101/600924

**Authors:** Ian AB Petersen, Kyle M Meyer, Brendan JM Bohannan

## Abstract

Bacterial communities are a major component of global diversity and are intimately involved in most terrestrial biogeochemical processes. Despite their importance, we know far less about the response of bacteria to human-induced environmental change than we do about other organisms. Understanding the response of organisms to land use change is especially pressing for tropical rainforests, which are being altered at a higher rate than any other ecosystem. Here, we conduct a meta-analysis of studies performed in each of the major tropical rainforest regions to ask whether there are consistent responses of belowground bacterial communities to the conversion of tropical rainforest to agriculture. Remarkably, we find common responses despite wide variation across studies in the types of agriculture practiced and the research methodology used to study land use change. These responses include changes in the relative abundance of phyla, most notably decreases in Acidobacteria and Proteobacteria and increases in Actinobacteria, Chloroflexi and Firmicutes. We also find that alpha diversity (at the scale of single soil cores), consistently increases with ecosystem conversion. These consistent responses suggest that, while there is great diversity in agricultural practices across the tropics, common features such as the use of slash-and-burn tactics have the potential to alter bacterial community composition and diversity belowground.

## 1 Introduction

There is growing recognition of the importance of understanding the response of bacterial communities to land use change. Soil bacteria mediate most biogeochemical cycles (Madsen, 2011) and comprise the majority of global biodiversity (Hug et al., 2016). When ecosystems undergo conversion, such as from forest to agriculture, biochemical cycles are often profoundly altered (DeFries, Field, Fung, Collatz, & Bounoua, 1999; Neill et al., 1997, 2005; Verchot, Davidson, Cattânio, & Ackerman, 2000) and the diversity of plant and animal communities can decrease, along with changes in species composition (Banks, Sandvik, & Keesecker, 2009; Bernard, Fjeldså, & Mohamed, 2009; Gardner et al., 2009; Gibson et al., 2011; Newbold et al., 2015; Perfecto & Snelling, 1995). No ecosystem type is experiencing conversion at a faster rate than tropical rainforests (Dirzo & Raven, 2003), which are declining primarily due to agricultural expansion (De Moraes, Volkoff, Cerri, & Bernoux, 1996). From 1980 to 2012, approximately 1.3 million km^2^ of tropical rainforest were cleared for agriculture, an area the size of France, Spain and the UK combined (Gibbs et al., 2010; Hansen et al., 2013).

Across the tropics, forest-to-agriculture conversion occurs through a similar mechanism, slash-and-burn, but differs in eventual agricultural characteristics. In slash-and-burn, valuable trees are removed for lumber and the remaining vegetation is burned to clear the land for agricultural use. These agricultural uses vary by region, with oil palm plantations particularly common in southeast Asia (Koh & Wilcove, 2008), beef cattle pastures or soybean plantation in the Amazon Basin (Morton et al., 2006) and small scale subsistence farming of plantain or manioc in the Congo Basin (Sunderlin et al., 2000; Zhang, Justice, Jiang, Brunner, & Wilkie, 2006). Duration of agricultural use also varies by region, from one-three years for plantations in the Congo (Zhang, Justice, & Desanker, 2002) to decades of continual use in southeast Asia and the Amazon (Rodrigues et al., 2013; Tripathi et al., 2016). While there is great diversity in agricultural characteristics across the tropics, consistent patterns in bacterial response should nonetheless emerge if community change is driven by common aspects of ecosystem conversion, such as slash-and-burn tactics.

There are a growing number of studies on the effect of land use change on belowground microbial communities in the tropics. We took advantage of this growing literature to ask whether there are commonalities across studies in the response of belowground microbial communities to ecosystem conversion. Specifically, we conducted a meta-analysis on the effects of forest to agriculture conversion on belowground bacterial communities and soil chemical properties across the world’s three largest tropical rainforests (the Amazon Basin, the Congo Basin and southeast Asia). Despite regional differences in agricultural practices, and a diversity of research methodologies across studies, we observed consistent changes in diversity and phylum level composition.

## 2 Methods

We conducted a literature search for studies that directly compare soil samples from primary tropical rainforest with proximal agricultural sites, analyze metabarcode or metagenome sequences and report at least one diversity metric, composition at the phylum level and soil chemical characteristics.

Following identification, means (*X*), standard deviations (SDs), and sample sizes (n) were extracted from the papers. If not reported, these statistics were derived from other metrics; standard error, 95% confidence interval, or diversity metrics and composition were derived and summarized from OTU tables using the vegan package in R 3.4.4. The meta-analysis was then conducted following Borenstein *et al*. (2009). Briefly, for each study a response ratio was determined for diversity, phylum-level relative abundance and soil chemical factors using the following equation

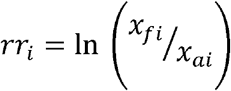

Where for study *i*, the means of forest and agricultural samples are represented respectively by *x_fi_* and *x_ai_*. To assess responses across all studies and a weighting value (*w*) was calculated.

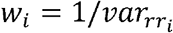

The summary response ratio (*rr_w_*) was then calculated by averaging individual response ratios and weighting values.

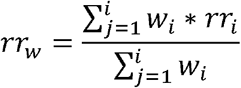

To determine if the summary response ratio was significantly different from zero, we then conducted two tailed t-tests (*α* = 0.05).

For phyla that we found to have a significant response, we then attempted to examine the consistency of response at finer taxonomic scales (order, class) using findings reported in the published papers or by deriving information from the underlying OTU tables.

## 3 Results

Our literature search identified seventeen studies (Supplementary Table 1), with representation from all three major rainforests (Figure 1). A diverse set of agriculture types were represented, including cattle pasture (predominantly in the Amazon), oil palm plantation (exclusively in southeast Asia) and banana/manioc plantation in the Congo basin (Supplementary Table 1). In addition to a diversity of study systems, a variety of sampling techniques were used; with soil cores collected along transects, quadrats or nested sampling schemes and sequencing of samples done individually or as pooled samples.

**Figure 1:**
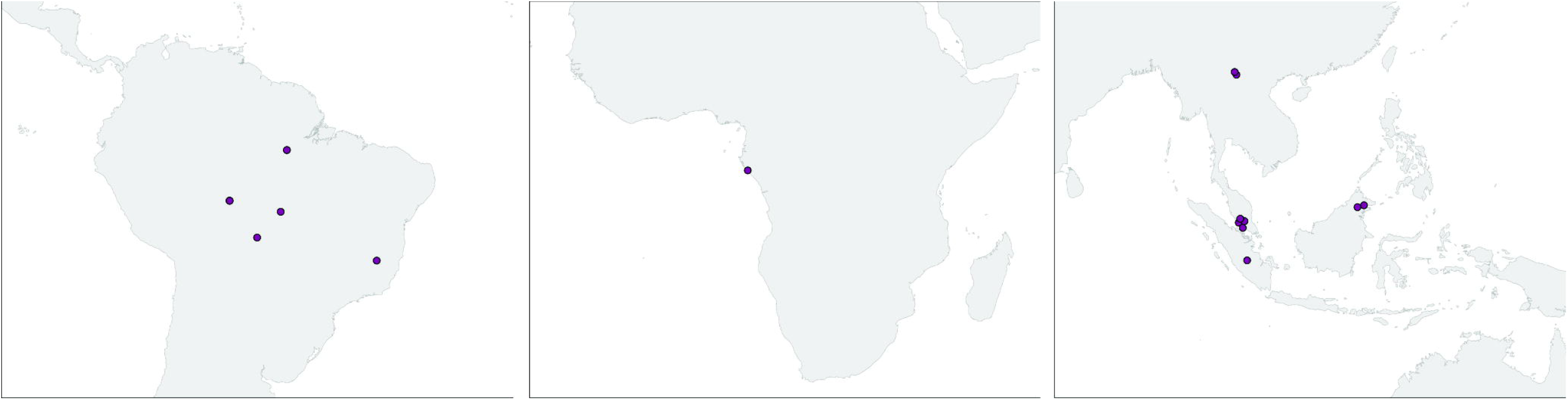
Study locations from the Amazon, Congo and SE Asia. Studies from the same site are represented by a single dot.

The V3 and/or the V4 region of 16S rRNA gene was sequenced in all studies using 454 pyrosequencing or the Illumina Hiseq or Miseq platforms. Three studies sequenced metagenomes, with one study solely analyzing metagenomic data and the other two pairing metagenomes with 16S metabarcoding (Supplementary Table 1). A wide variety of diversity metrics were used to quantify alpha diversity across studies. The Shannon index was the most commonly reported, followed by richness, with other metrics (Simpson, Faiths PD, Hills effective species number) included in only a few studies. There was not a single diversity metric reported in every study, but since response ratio is a unitless metric, differing metrics were considered together as alpha diversity.

### 3.1 Diversity

Alpha diversity results were reported in all but one study. In these studies, the majority, (9) reported increases in alpha diversity with conversion (p>0.001), six reported no change, and one study reported a decrease (Figure 2). In contrast, beta diversity was not reported in a majority of studies. Among the eight studies that reported beta diversity, a wide variety of sampling and statistical approaches were used, and the results were highly variable. Significant decreases in beta diversity following conversion were reported in three studies, while two studies reported significant increases, and the remainder reporting no consistent response.

**Figure 2:**
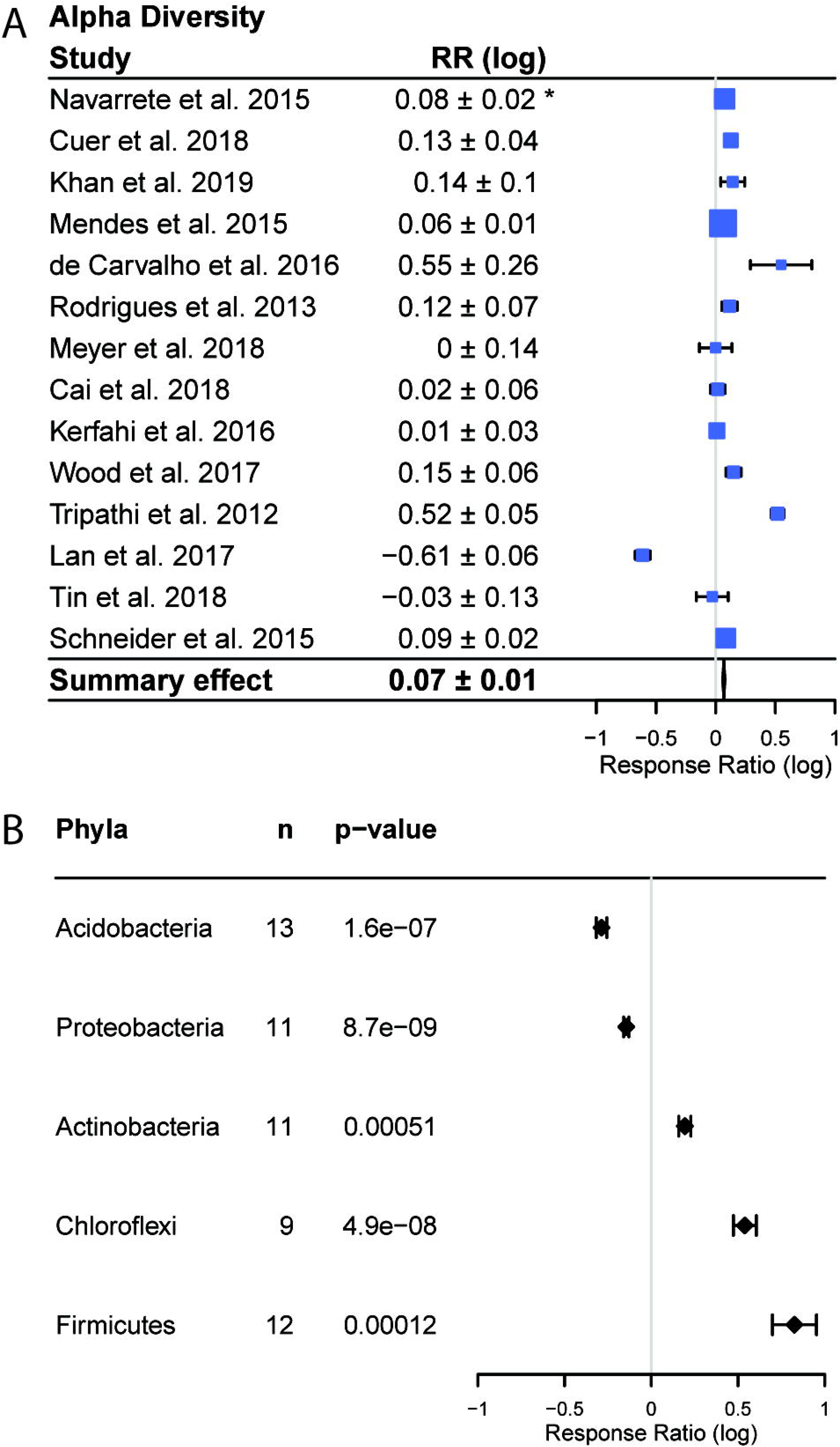
Effects of forest to agriculture conversion on bacterial diversity and composition. (a) Changes in alpha diversity measured as the response ratio (RR). Positive values indicate higher diversity in agricultural areas and negative values indicate higher diversity in forest areas. The bars represent the 95% confidence intervals (CIs) and box size corresponds to study weight in the summary effect size. Summary response ratio was significantly different from zero (two tailed t-tests, p < 0.001). (b) Changes in abundance of phyla reported in a majority of studies, including the number of studies reporting each phylum out of 16 studies. * Confidence interval (95%) of the response ratio.

### 3.2 Composition

Bacterial community composition varied across studies, but members of five phyla (Actinobacteria, Firmicutes, Chloroflexi, Acidobacteria, Proteobacteria) were reported in a majority of studies (Figure 2). We observed consistent responses to land use change in each of these five phyla (all p<0.01). The relative abundances of Actinobacteria, Chloroflexi and Firmicutes increased with conversion, while the relative abundance of the remaining two phyla, Acidobacteria and Proteobacteria, decreased (Figure 2). In addition to having overall significant effects, these responses were also remarkably universal across studies (Figure 3). Only two studies reported any changes in abundance that were opposite to the summary effect; Firmicutes were reported to decrease in Cuer *et al*. (2018), and Schneider *et al*. (2015) reported a decrease in Actinobacteria and increase in Acidobacteria with conversion (Figure 3).

**Figure 3:**
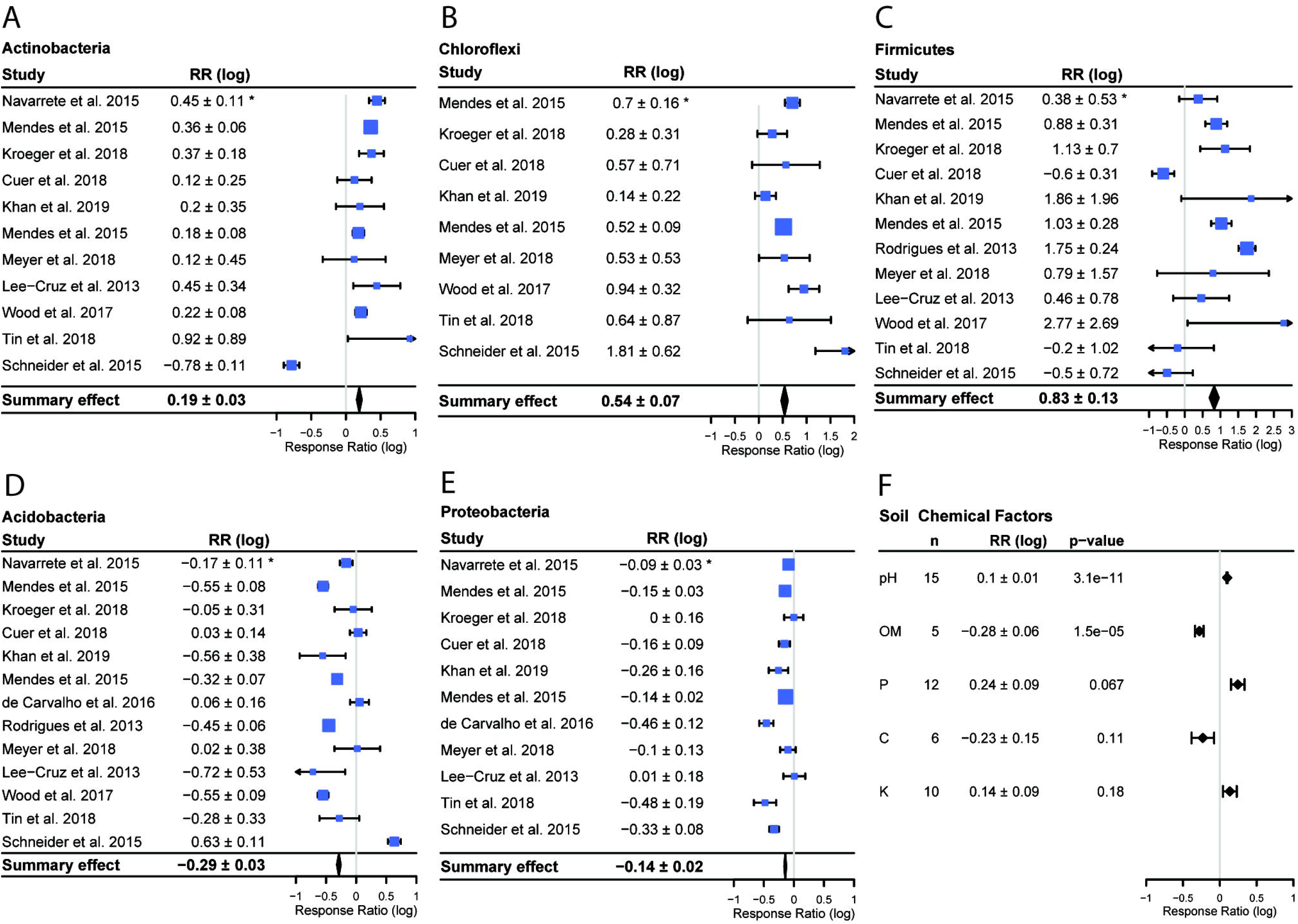
Forest plot for each phylum with a significant summary effect (all p < 0.001) (a-e) and summary effects for soil chemical data (f). Bars denote 95% confidence intervals, boxes are centered on means and box size corresponds to study weight for calculating summary effect. * Confidence interval (95%) of the response ratio.

Within the five phyla exhibiting consistent responses, we looked for consistencies at finer taxonomic levels, either by analyzing available OTU tables or by compiling findings reported in the papers. Such data were rare among the studies; only three reported responses of at least some classes or orders, and four provided OTU tables. Within these seven studies, we found no consistent responses at the class or order level within the phyla that exhibited consistent responses. Thus, while we have no evidence for common responses at finer taxonomic levels, it should be noted that this hypothesis was tested using far fewer studies than used for testing phylum level hypotheses.

### 3.3 Soil Chemical Factors

The only soil chemical factor that was both reported in a majority of studies and varied significantly with conversion was pH (p<0.001), which increased following conversion (Figure 3). The other two soil chemical factors reported in a majority of studies, potassium (K) and phosphorus (P), did not vary significantly with conversion. Organic matter decreased significantly with conversion, but it was only reported in 5 of the studies.

## 4 Discussion

Tropical rainforests harbor greater biodiversity than any other terrestrial ecosystem and are important contributors to global biogeochemical processes (Gardner et al., 2009). These ecosystems are being lost rapidly, primarily from land use conversion for agriculture (Gibbs et al., 2010; Hansen et al., 2013). The effects of land use change have only recently been studied in bacteria, which comprise a majority of global biodiversity and which mediate most biogeochemical cycles (Hug et al., 2016; Madsen, 2011). By conducting a meta-analysis, we found responses in the composition and diversity of soil bacterial communities to land use change spanning the world’s tropical zone.

### 4.1 Diversity

The increase in bacterial alpha diversity following conversion to agriculture differs sharply with commonly reported decreases in the diversity of plants (Ibrahim et al., 2003), animals (Bierregaard, 2001; Brook, Sodhl, & Ng, 2003; Gardner et al., 2009) and fungi (Cai, Zhang, Yang, & Wang, 2018; Kerfahi, Tripathi, Dong, Go, & Adams, 2016; Lan et al., 2017; McGuire et al., 2015; Mueller et al., 2014). This difference may result from ecological differences between bacteria and eukaryotes; for example, bacterial diversity may respond to changes in soil chemical properties, rather than being directly affected by land use change. We found that pH consistently increased with land use change across the studies we surveyed, and others have observed that pH can be correlated with bacterial richness (Fierer & Jackson, 2006; Lauber, Hamady, Knight, & Fierer, 2009). Aboveground productivity could also drive this trend. Agricultural management often promotes high productivity, which could increase soil labile carbon, driving increased bacterial diversity (Dias-Filho, Davidson, & Carvalho, 2001). Alternatively, the apparent increase in bacterial diversity could be an artifact of sampling. Increased alpha diversity at the level of a soil core may not represent landscape-level responses (*i.e*. gamma diversity). Bacteria may actually experience a landscape-level loss of diversity, similar to macro-organisms, if decreases in beta diversity offset or overwhelm increases in alpha diversity (as in Rodrigues *et al*. 2013). If this is the case, the apparent differences may be more methodological than ecological. Unfortunately, we cannot address this hypothesis because beta diversity was not often reported in the studies we surveyed and when measured was assessed with a wide variety of approaches. Consequently, we found that the results of beta diversity assessments were mixed. The difference in responses of eukaryotic and bacterial diversity to land use change should be investigated further to determine if this results from legitimate ecological differences or differences due to experimental design.

### 4.2 Composition

The commonality of community composition responses at the phylum level is striking and could suggest ecological trait conservation at the phylum level. Land use change is commonly associated with numerous changes to environmental factors (e.g. pH, P, K etc.) and biotic factors (mixing of species, introduction of new species, altered dispersal barriers, etc.). The response of an organism to these changes could depend on traits such as pH tolerance, life history strategies, stress tolerance, or dispersal ability. While the response to land use change is not a direct measurement of a particular trait, it could be thought of as the encapsulation of many individual traits, similar to a polygenetic phenotype. Because traits, especially complex traits, tend to be conserved phylogenetically (A. C. Martiny, Treseder, & Pusch, 2012; J. B. H. Martiny, Jones, Lennon, & Martiny, 2015), responses that are driven by complex combinations of traits should be conserved. We indeed find such conservation here, with consistent declines in Acidobacteria and Proteobacteria, and increases in Actinobacteria, Firmicutes and Chloroflexi. These common responses in composition support the notion of phylum-level conservation of traits that mediate land use change responses.

### 4.3 Functional consequences

Shifts in the diversity and composition of soil bacterial communities may have important implications for ecosystem function. Higher diversity levels have been associated with increased resistance, resilience, and/or stability of ecosystem function for a variety of aboveground ecosystems (Flynn, Mirotchnick, Jain, Palmer, & Naeem, 2011; Hooper et al., 2012; Tilman, Reich, & Isbell, 2012). The generalizability of this trend, however, is an ongoing debate (Cadotte, Carscadden, & Mirotchnick, 2011; Srivastava & Vellend, 2005). Community composition has also been reported to affect ecosystem function (Panke-Buisse, Poole, Goodrich, Ley, & Kao-Kniffin, 2014; Swenson, Arendt, & Wilson, 2000; Swenson, Wilson, & Elias, 2000). A recent meta-study found that incorporating microbial composition data with environmental data improves predictions of ecosystem process rates in 53% of cases (Graham et al., 2016), suggesting that widespread changes to bacterial diversity and composition could impact the functioning of ecosystems. However, we did not explicitly test this hypothesis in our study.

### 4.4 Recommendations for future research

The apparent commonality of bacterial responses warrants a more detailed examination, and we suggest several paths forward. First, there is a need to increase the geographic representation of key areas, especially the Congo River Basin. Our comparison is limited by the number of studies fitting our criteria. Tropical ecosystems in general have received considerably less attention from microbial ecologists than temperate areas (Pajares, Bohannan, & Souza, 2016). Encouragingly, bacterial responses to land use change is an emerging area of study, with every study we compared published since 2012. Increasing the number of studies, particularly in the Congo Basin, will allow more rigorous testing of patterns we report, and would provide sufficient power to assess the importance of other characteristics, such as agricultural form and duration of agricultural use, on bacterial responses.

Second, inclusion of other soil organisms such as fungi, nematodes and microarthropods will allow for a more complete understanding of how soil ecosystems are affected by land use change (as in Kerfahi *et al*. 2016). Furthermore, simultaneous sampling of soil biota would allow the direct comparison of responses across large taxonomic groups; this could reveal fundamentally different responses to land use change across these groups. For example, while only five studies of tropical land use change to date have focused on fungi, four reported decreases in alpha diversity with conversion, suggesting that fungi may respond differently to ecosystem conversion than bacteria.

Finally, as more studies are conducted, increased standardization should be adopted for sampling protocols and reporting of findings. The studies assessed here span a breadth of protocols for sampling and a variety of metrics to assess diversity and composition. The generality of findings across this breadth points to the strength of the trends we observed. However, we were hampered in our ability to assess consistent changes in class level abundances and beta diversity due to low levels of reporting and by limited overlap in sampling protocols and analyses. Specifically, we suggest reporting GPS coordinates for every sample, reporting common diversity metrics (Shannon, richness etc.), making OTU tables and raw sequences available, and we recommend not pooling soil samples (so that spatially explicit analyses are possible). A more standardized set of collection and assessment methods will allow for a more conclusive and thorough analysis of the relationship between land use change and community attributes. If the above steps are taken, we believe it will lead to a much richer understanding of how land use change affects tropical soil communities.

### 4.5 Conclusions

We compared studies spanning a global longitudinal range, with representation from the three largest rainforests, each of which harbors vast biodiversity and high levels of endemism. The studies we compared varied in the primary form of agriculture, duration of agricultural use and research methodology. Despite this variation, we identified consistent shifts in the diversity and composition of bacterial communities in response to the conversion of tropical rainforests to agriculture. Our findings suggest that land use conversion can have predictable effects at a pan-tropical level. With tropical rainforests undergoing dramatic and rapid reductions, it is important to consider the ramifications such changes will have on belowground soil communities and the ecosystem processes they mediate.

#### Contribution to the field

Interest in studying the response of soil bacterial communities to forest-to-agriculture conversion in the tropics has increased dramatically since 2012. However there has not yet been an effort to quantitatively ask whether responses are consistent. To provide this assessment, we conducted a meta-analysis on a set of studies with pan-tropical representation and wide variation in agricultural characteristics and research methodology. Despite this variation, we identified consistent changes in the diversity and composition of belowground bacterial communities. These consistent responses suggest that, while there is great diversity in agricultural practices across the tropics, common features such as the use of slash-and-burn tactics have the potential to alter bacterial community composition and diversity belowground. With tropical rainforests undergoing dramatic and rapid reductions, it is important to consider the ramifications such changes will have on belowground soil communities and the ecosystem processes they mediate.

## Acknowledgements

We thank the authors of the studies we analyzed, without which this meta-analysis would not be possible. This work was supported by the National Science Foundation – Dimensions of Biodiversity (DEB 14422214).

## Author Contributions

IP, KM and BJMB designed the study. IP conducted the literature search, meta-analysis and wrote the manuscript. KM, BJMB and IP contributed to manuscript revisions, and read and approved the final version.

---

**Supplementary Table 1:**
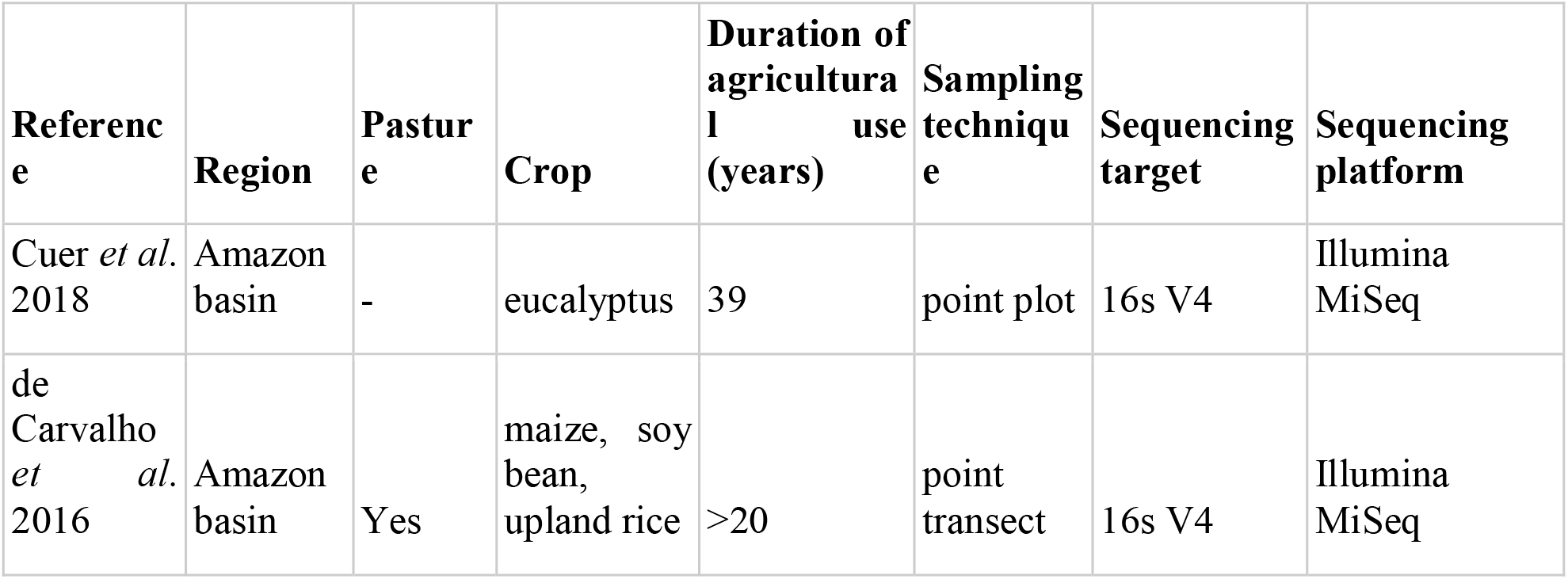
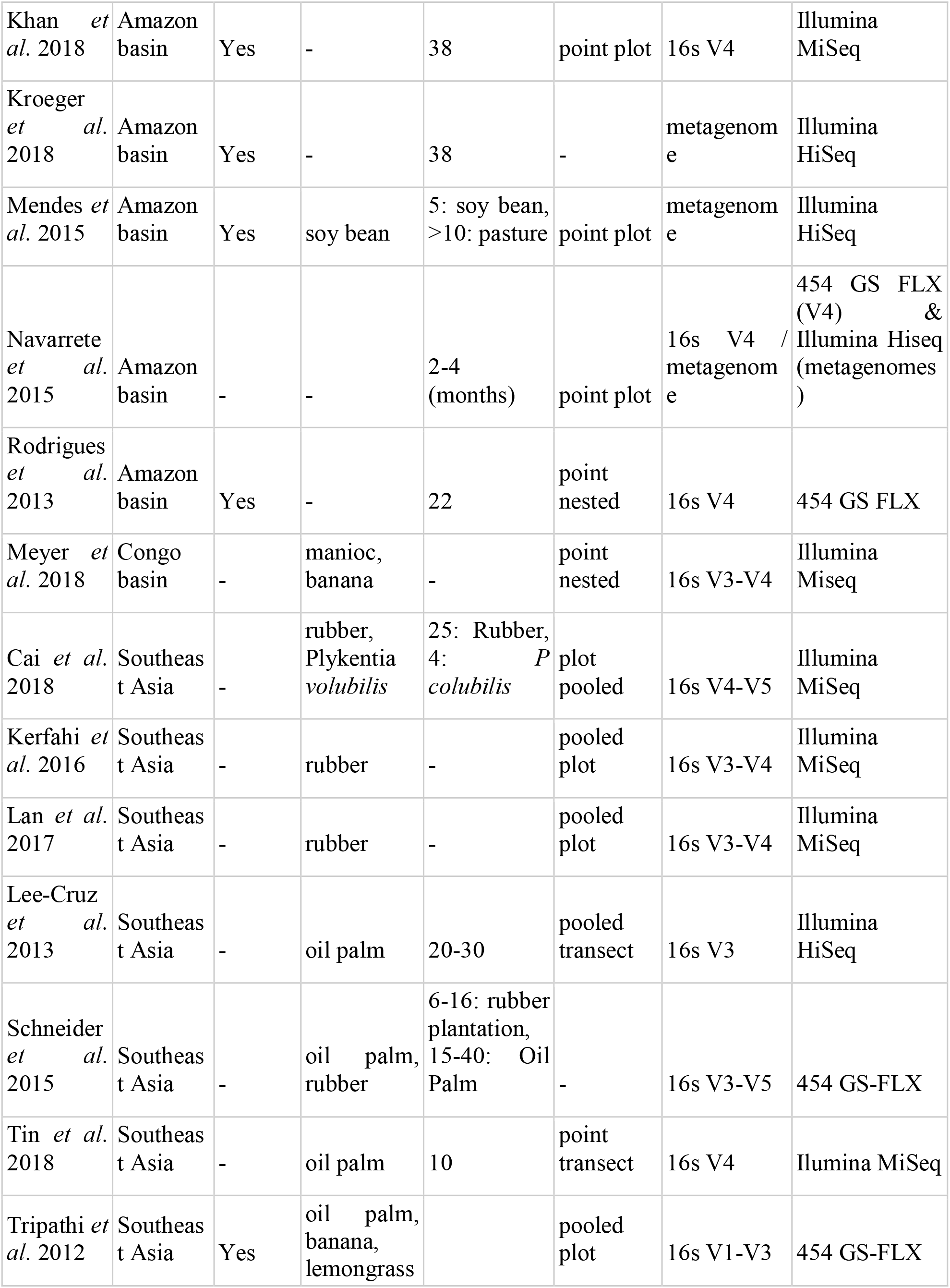
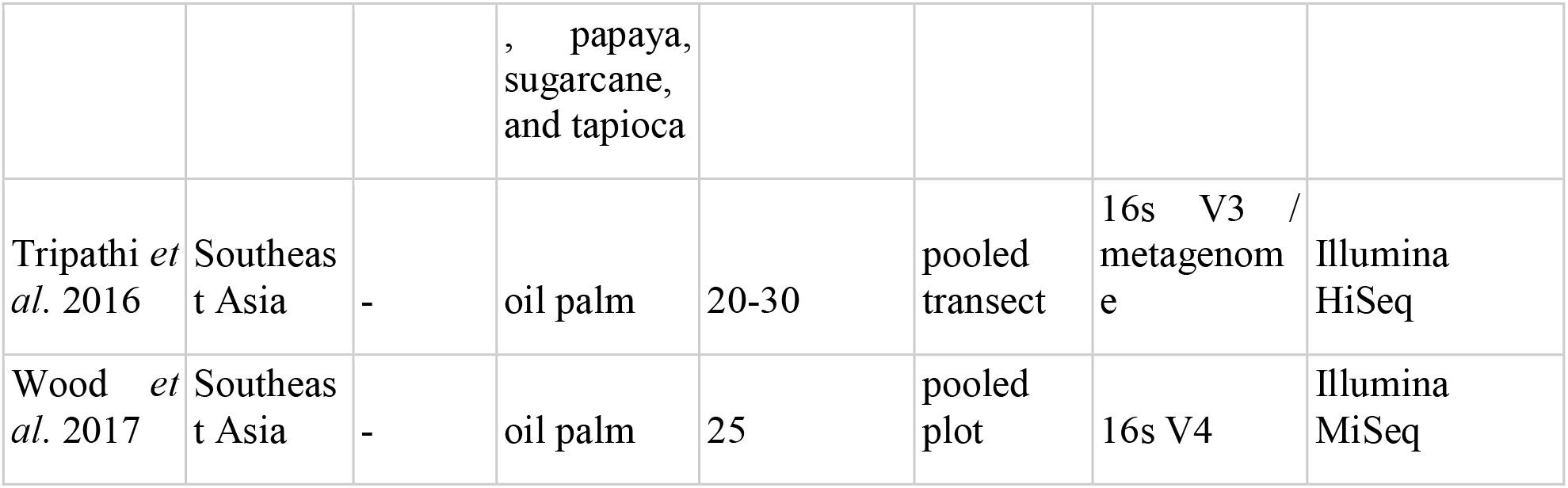
Summary of included studies.

| Reference | Region | Pasture | Crop | Duration of agriculture use (years) | Sampling technique | Sequencing target | Sequencing platform |
| --- | --- | --- | --- | --- | --- | --- | --- |
| Cuer *et al.* 2018 | Amazon basin | - | eucalyptus | 39 | point plot | 16s V4 | Illumina MiSeq |
| de Carvalho *et al.* 2016 | Amazon basin | Yes | maize, soy bean, upland rice | >20 | point transect | 16s V4 | Illumina MiSeq |
| Pan-tropical bacterial responses to deforestation |  |  |  |  |  |  |  |
| Khan *et al.* 2018 | Amazon basin | Yes | - | 38 | point plot | 16s V4 | Illumina MiSeq |
| Kroeger *et al.* 2018 | Amazon basin | Yes | - | 38 | - | metagenome | Illumina HiSeq |
| Mendes *et al.* 2015 | Amazon basin | Yes | soy bean | 5: soy bean, >10: pasture | point plot | metagenome | Illumina HiSeq |
| Navarrete *et al.* 2015 | Amazon basin | - | - | 2-4 (months) | point plot | 16s V4 / metagenome | 454 GS FLX (V4) & Illumina HiSeq (metagenomes) |
| Rodrigues *et al.* 2013 | Amazon basin | Yes | - | 22 | point nested | 16s V4 | 454 GS FLX |
| Meyer *et al.* 2018 | Congo basin | - | manioc, banana | - | point nested | 16s V3-V4 | Illumina Miseq |
| Cai *et al.* 2018 | Southeast Asia | - | rubber, *Plykrentia volubilis* | 25: Rubber, 4: *P. colubilis* | plot pooled | 16s V4-V5 | Illumina MiSeq |
| Kerfahi *et al.* 2016 | Southeast Asia | - | rubber | - | pooled plot | 16s V3-V4 | Illumina MiSeq |
| Lan *et al.* 2017 | Southeast Asia | - | rubber | - | pooled plot | 16s V3-V4 | Illumina MiSeq |
| Lee-Cruz *et al.* 2013 | Southeast Asia | - | oil palm | 20-30 | pooled transect | 16s V3 | Illumina HiSeq |
| Schneider *et al.* 2015 | Southeast Asia | - | oil palm, rubber | 6-16: rubber plantation, 15-40: Oil Palm | - | 16s V3-V5 | 454 GS-FLX |
| Tin *et al.* 2018 | Southeast Asia | - | oil palm | 10 | point transect | 16s V4 | Illumina MiSeq |
| Tripathi *et al.* 2012 | Southeast Asia | Yes | oil palm, banana, lemongrass |  | pooled plot | 16s V1-V3 | 454 GS-FLX |
|  |  |  | , papaya,<br>sugarcane,<br>and tapioca |  |  |  |  |
| Tripathi *et al.* 2016 | Southeast Asia | - | oil palm | 20-30 | pooled transect | 16s V3 / metagenome | Illumina HiSeq |
| Wood *et al.* 2017 | Southeast Asia | - | oil palm | 25 | pooled plot | 16s V4 | Illumina MiSeq |

## References

Banks, J. E., Sandvik, P., & Keesecker, L. (2009). Beetle (Coleoptera) and spider (Araneae) diversity in a mosaic of farmland, edge, and tropical forest habitats in western Costa Rica. The Pan-Pacific Entomologist, 83(2), 152–160. http://doi.org/10.3956/0031-0603-83.2.152

Bernard, H., Fjeldså, J., & Mohamed, M. (2009). A Case Study on the Effects of Disturbance and Conversion of Tropical Lowland Rain Forest on the Non-Volant Small Mammals in North Borneo: Management Implications. Mammal Study, 34(2), 85–96. http://doi.org/10.3106/041.034.0204

Bierregaard, R. O. (2001). Lessons from Amazonia□; the ecology and conservation of a fragmented forest. Yale University Press. Retrieved from https://books.google.com/books?hl=en&lr=&id=s3YQx894sqIC&oi=fnd&pg=PP13&dq=Lessons+from+Ama-+zonia:+The+Ecology+and+Conservation+of+a+Fragmented+Forest&ots=9tgbSWewiU&sig=e8Gdu8HfgWnYv6XesStDq5evtEw#v=onepage&q=Lessons from Ama- zonia%3A The Ecology and C

Borenstein, M., Hedges, L. V, Higgins, J. P. T., & Rothstein, H. R. (2009). Introduction to Meta-Analysis. Handbook of Set Theoretic Topology. Retrieved from www.wiley.com.

Brook, B. W., Sodhl, N. S., & Ng, P. K. L. (2003). Catastrophic extinctions follow deforestation in Singapore. Nature, 424(6947), 420–423. http://doi.org/10.1038/nature01795

Cadotte, M. W., Carscadden, K., & Mirotchnick, N. (2011). Beyond species: functional diversity and the maintenance of ecological processes and services. Journal of Applied Ecology, 48(5), 1079–1087. http://doi.org/10.1111/j.1365-2664.2011.02048.x

Cai, Z., Zhang, Y., Yang, C., & Wang, S. (2018). Land-use type strongly shapes community composition, but not always diversity of soil microbes in tropical China. Catena, 165, 369–380. http://doi.org/10.1016/J.CATENA.2018.02.018

Cuer, C. A., Rodrigues, R. de A. R., Balieiro, F. C., Jesus, J., Silva, E. P., Alves, B. J. R., & Rachid, C. T. C. C. (2018). Short-term effect of Eucalyptus plantations on soil microbial communities and soil-atmosphere methane and nitrous oxide exchange. Scientific Reports, 8(1), 15133. http://doi.org/10.1038/s41598-018-33594-6

De Moraes, J. F. L., Volkoff, B., Cerri, C. C., & Bernoux, M. (1996). Soil properties under Amazon forest and changes due to pasture installation in Rondônia, Brazil. Geoderma, 70(1), 63–81. http://doi.org/10.1016/0016-7061(95)00072-0

DeFries, R. S., Field, C. R., Fung, I., Collatz, G. J., & Bounoua, L. (1999). Combining satellite data and bioegeochemical models to estimate global effects of human-induced land cover change on carbon emissions and primary productivity. Global Biogeochemical Cycles, 13(3), 803–815.

Dias-Filho, M., Davidson, E., & Carvalho, C. (2001). Linking biogeochemical cycles to cattle pasture management and sustainability in the Amazon Basin. The Biogeochemistry of the Amazon Basin, (July 2015), 86–101. Retrieved from https://books.google.com/books?hl=en&lr=&id=3JLmCwAAQBAJ&oi=fnd&pg=PA84&dq=)+Linking+biogeochemical+cycles+to+cattle+pasture+management+and+sustainability+in+the+Amazon+Basin.+The+Biogeochemistry+of+the+Amazon+Basin+(ed.+by+M.E.+McClain,+R.L.+Victoria,+an

Dirzo, R., & Raven, P. H. (2003). Global State of Biodiversity and Loss. Annual Review of Environment and Resources, 28(1), 137–167. http://doi.org/10.1146/annurev.energy.28.050302.105532

Fierer, N., & Jackson, R. (2006). The diversity and biogeography of soil bacterial communities. Proceedings of the National Academy of Sciences, 103(3), 626–631. http://doi.org/10.1073/pnas.95.12.6578

Flynn, D. F. B., Mirotchnick, N., Jain, M., Palmer, M. I., & Naeem, S. (2011). Functional and phylogenetic diversity as predictors of biodiversity–ecosystem-function relationships. Ecology, 92(8), 1573–1581. http://doi.org/10.1890/10-1245.1

Gardner, T. A., Barlow, J., Chazdon, R., Ewers, R. M., Harvey, C. A., Peres, C. A., & Sodhi, N. S. (2009). Prospects for tropical forest biodiversity in a human-modified world. Ecology Letters, 12(6), 561–582. http://doi.org/10.1111/j.1461-0248.2009.01294.x

Gibbs, H. K., Ruesch, A. S., Achard, F., Clayton, M. K., Holmgren, P., Ramankutty, N., & Foley, J. A. (2010). Tropical forests were the primary sources of new agricultural land in the 1980s and 1990s. Proceedings of the National Academy of Sciences, 107(38), 16732–16737. http://doi.org/10.1073/pnas.0910275107

Gibson, L., Lee, T. M., Koh, L. P., Brook, B. W., Gardner, T. A., Barlow, J., … Sodhi, N. S. (2011). Primary forests are irreplaceable for sustaining tropical biodiversity. Nature, 478(7369), 378–381. http://doi.org/10.1038/nature10425

Graham, E. B., Knelman, J. E., Schindlbacher, A., Siciliano, S., Breulmann, M., Yannarell, A., … Nemergut, D. R. (2016). Microbes as Engines of Ecosystem Function: When Does Community Structure Enhance Predictions of Ecosystem Processes? Frontiers in Microbiology, 7(FEB), 1–10. http://doi.org/10.3389/fmicb.2016.00214

Hansen, M. C. C., Potapov, P. V, Moore, R., Hancher, M., Turubanova, S. A. a, Tyukavina, A., … Townshend, J. R. G. R. G. (2013). High-resolution global maps of 21st-century forest cover change. Science, 342(November), 850–854. http://doi.org/10.1126/science.1244693

Hooper, D. U., Adair, E. C., Cardinale, B. J., Byrnes, J. E. K., Hungate, B. A., Matulich, K. L., … O’Connor, M. I. (2012). A global synthesis reveals biodiversity loss as a major driver of ecosystem change. Nature, 486(7401), 105–108. http://doi.org/10.1038/nature11118

Hug, L. A., Baker, B. J., Anantharaman, K., Brown, C. T., Probst, A. J., Castelle, C. J., … Banfield, J. F. (2016). A new view of the tree of life. Nature Microbiology, 1(5), 16048. http://doi.org/10.1038/nmicrobiol.2016.48

Ibrahim, A. Bin, Wee, Y. C., Turner, I. M., Corlett, R. T., Chew, P. T., & Tan, H. T. W. (2003). A Study of Plant Species Extinction in Singapore: Lessons for the Conservation of Tropical Biodiversity. Conservation Biology, 8(3), 705–712. http://doi.org/10.1046/j.1523-1739.1994.08030705.x

Kerfahi, D., Tripathi, B. M., Dong, K., Go, R., & Adams, J. M. (2016). Rainforest Conversion to Rubber Plantation May Not Result in Lower Soil Diversity of Bacteria, Fungi, and Nematodes. Microbial Ecology, 72(2), 359–371. http://doi.org/10.1007/s00248-016-0790-0

Koh, L. P., & Wilcove, D. S. (2008). Is oil palm agriculture really destroying tropical biodiversity? Conservation Letters, 1(2), 60–64. http://doi.org/10.1111/j.1755-263X.2008.00011.x

Lan, G., Li, Y., Jatoi, M. T., Tan, Z., Wu, Z., & Xie, G. (2017). Change in Soil Microbial Community Compositions and Diversity Following the Conversion of Tropical Forest to Rubber Plantations in Xishuangbanan, Southwest China. Tropical Conservation Science, 10, 1–14. http://doi.org/10.1177/1940082917733230

Lauber, C. L., Hamady, M., Knight, R., & Fierer, N. (2009). Pyrosequencing-based assessment of soil pH as a predictor of soil bacterial community structure at the continental scale. Applied and Environmental Microbiology, 75(15), 5111–5120. http://doi.org/10.1128/AEM.00335-09

Madsen, E. L. (2011). Microorganisms and their roles in fundamental biogeochemical cycles. Current Opinion in Biotechnology, 22, 456–464. http://doi.org/10.1016/j.copbio.2011.01.008

Martiny, A. C., Treseder, K., & Pusch, G. (2012). Phylogenetic conservatism of functional traits in microorganisms. The ISME Journal, 7(10), 830–838. http://doi.org/10.1038/ismej.2012.160

Martiny, J. B. H., Jones, S. E., Lennon, J. T., & Martiny, A. C. (2015). Microbiomes in light of traits: A phylogenetic perspective. Science (New York, N.Y.), 350(6261), aac9323. http://doi.org/10.1126/science.aac9323

McGuire, K. L., D’Angelo, H., Brearley, F. Q., Gedallovich, S. M., Babar, N., Yang, N., … Fierer, N. (2015). Responses of Soil Fungi to Logging and Oil Palm Agriculture in Southeast Asian Tropical Forests. Microbial Ecology, 69(4), 733–747. http://doi.org/10.1007/s00248-014-0468-4

Morton, D. C., DeFries, R. S., Shimabukuro, Y. E., Anderson, L. O., Arai, E., del Bon Espirito-Santo, F., … Morisette, J. (2006). Cropland expansion changes deforestation dynamics in the southern Brazilian Amazon. Proceedings of the National Academy of Sciences, 103(39), 14637–14641. http://doi.org/10.1073/pnas.0606377103

Mueller, R. C., Paula, F. S., Mirza, B. S., Rodrigues, J. L. M., Nüsslein, K., & Bohannan, B. J. M. (2014). Links between plant and fungal communities across a deforestation chronosequence in the Amazon rainforest. ISME Journal, 8(7), 1548–1550. http://doi.org/10.1038/ismej.2013.253

Neill, C., Piccolo, M. C., Cerri, C. C., Steudler, P. A., Melillo, J. M., & Brito, M. (1997). Net nitrogen mineralization and net nitrification rates in soils following deforestation for pasture across the southwestern Brazilian Amazon Basin landscape. Oecologia, 110(2), 243–252. http://doi.org/10.1007/s004420050157

Neill, C., Steudler, P. A., Garcia-Montiel, D. C., Melillo, J. M., Feigl, B. J., Piccolo, M. C., & Cerri, C. C. (2005). Rates and controls of nitrous oxide and nitric oxide emissions following conversion of forest to pasture in Rondônia. Nutrient Cycling in Agroecosystems, 71(1), 1–15. http://doi.org/10.1007/s10705-004-0378-9

Newbold, T., Hudson, L. N., Hill, S. L. L., Contu, S., Lysenko, I., Senior, R. A., … Purvis, A. (2015). Global effects of land use on local terrestrial biodiversity. Nature, 520(7545), 45–50. http://doi.org/10.1038/nature14324

Pajares, S., Bohannan, B. J. M., & Souza, V. (2016). Editorial: The Role of Microbial Communities in Tropical Ecosystems. Frontiers in Microbiology, 7, 1805. http://doi.org/10.3389/fmicb.2016.01805

Panke-Buisse, K., Poole, A. C., Goodrich, J. K., Ley, R. E., & Kao-Kniffin, J. (2014). Selection on soil microbiomes reveals reproducible impacts on plant function. The ISME Journal, 9(4), 980–989. http://doi.org/10.1038/ismej.2014.196

Perfecto, I., & Snelling, R. (1995). Biodiversity and the Transformation of a Tropical Agroecosystem: Ants in Coffee Plantations. Ecological Applications, 5(4), 1084–1097. http://doi.org/10.2307/2269356

Rodrigues, J. L. M., Pellizari, V. H., Mueller, R., Baek, K., Jesus, E. D. C., Paula, F. S., … Nüsslein, K. (2013). Conversion of the Amazon rainforest to agriculture results in biotic homogenization of soil bacterial communities. Proceedings of the National Academy of Sciences of the United States of America, 110(3), 988–993. http://doi.org/10.1073/pnas.1220608110

Schneider, D., Engelhaupt, M., Allen, K., Kurniawan, S., Krashevska, V., Heinemann, M., … Daniel, R. (2015). Impact of lowland rainforest transformation on diversity and composition of soil prokaryotic communities in sumatra (Indonesia). Frontiers in Microbiology, 6(DEC), 1339. http://doi.org/10.3389/fmicb.2015.01339

Srivastava, D. S., & Vellend, M. (2005). Biodiversity-Ecosystem Function Research: Is It Relevant to Conservation? Annual Review of Ecology, Evolution, and Systematics, 36(1), 267–294. http://doi.org/10.1146/annurev.ecolsys.36.102003.152636

Sunderlin, W. D., Ndoye, O., Bikie, H., Laporte, N., Mertens, B., & Pokam, J. (2000). Economic crisis, small-scale agriculture, and forest cover change in southern Cameroon. Environmental Conservation, 27(3), 284–290. http://doi.org/10.1017/s0376892900000321

Swenson, W., Arendt, J., & Wilson, D. S. (2000). Artificial selection of microbial ecosystems for 3-chloroaniline biodegradation. Environmental Microbiology, 2(5), 564–571. http://doi.org/10.1046/j.1462-2920.2000.00140.x

Swenson, W., Wilson, D. S., & Elias, R. (2000). Artificial ecosystem selection. Proceedings of the National Academy of Sciences of the United States of America, 97(16), 9110–4. http://doi.org/10.1073/pnas.150237597

Tilman, D., Reich, P. B., & Isbell, F. (2012). Biodiversity impacts ecosystem productivity as much as resources, disturbance, or herbivory. Proceedings of the National Academy of Sciences of the United States of America, 109(26), 10394–7. http://doi.org/10.1073/pnas.1208240109

Tripathi, B. M., Edwards, D. P., Mendes, L. W., Kim, M., Dong, K., Kim, H., & Adams, J. M. (2016). The Impact of tropical forest logging and oil palm agriculture on the soil microbiome. Molecular Ecology, 25(10), 2244–2257. http://doi.org/10.1111/mec.13620

Verchot, L. V., Davidson, E. A., Cattânio, J. H., & Ackerman, I. L. (2000). Land-use change and biogeochemical controls of methane fluxes in soils of eastern Amazonia. Ecosystems, 3(1), 41–56. http://doi.org/10.1007/s100210000009

Zhang, Q., Justice, C. O., & Desanker, P. V. (2002). Impacts of simulated shifting cultivation on deforestation and the carbon stocks of the forests of central Africa. Agriculture, Ecosystems and Environment, 90(2), 203–209. http://doi.org/10.1016/S0167-8809(01)00332-2

Zhang, Q., Justice, C. O., Jiang, M., Brunner, J., & Wilkie, D. S. (2006). A gis-based assessment on the vulnerability and future extent of the tropical forests of the Congo Basin. Environmental Monitoring and Assessment, 114(1–3), 107–121. http://doi.org/10.1007/s10661-006-2015-3

## References

de Carvalho, T. S., Jesus, E. da C., Barlow, J., Gardner, T. A., Soares, I. C., Tiedje, J. M., & Moreira, F. M. de S. (2016). Land use intensification in the humid tropics increased both alpha and beta diversity of soil bacteria. Ecology, 97(10), 2760–2771. http://doi.org/10.1002/ecy.1513

Khan, M. A. W., Bohannan, B. J. M., Nüsslein, K., Tiedje, J. M., Tringe, S. G., Parlade, E., … Rodrigues, J. L. M. (2018). Deforestation Impacts Network Co-occurrence Patterns of Microbial Communities in Amazon Soils. FEMS Microbiology Ecology, (April 2018), 1–12. http://doi.org/10.1093/femsec/fiy230

Kroeger, M. E., Delmont, T. O., Eren, A. M., Meyer, K. M., Guo, J., Khan, K., … Nüsslein, K. (2018). New biological insights into how deforestation in amazonia affects soil microbial communities using metagenomics and metagenome-assembled genomes. Frontiers in Microbiology, 9(JUL), 1635. http://doi.org/10.3389/fmicb.2018.01635

Lee-Cruz, L., Edwards, D. P., Tripathi, B. M., & Adams, J. M. (2013). Impact of logging and forest conversion to oil palm on soil bacterial communities in Borneo. SUPP. Applied and Environmental Microbiology, 79, 7290–7297. http://doi.org/10.1128/AEM.02541-13

Mendes, L. W., Tsai, S. M., Navarrete, A. A., de Hollander, M., van Veen, J. A., & Kuramae, E. E. (2015). Soil-Borne Microbiome: Linking Diversity to Function. Microbial Ecology, 70(1), 255–265. http://doi.org/10.1007/s00248-014-0559-2

Meyer, K. M., Petersen, I. A. B., Tobi, E., Korte, L., & Bohannan, B. (2018). Use of RNA and DNA to Identify Mechanisms of Microbial Community Homogenization. BioRxiv, 496679. http://doi.org/10.1101/496679

Navarrete, A. A., Tsai, S. M., Mendes, L. W., Faust, K., De Hollander, M., Cassman, N. A., … Kuramae, E. E. (2015). Soil microbiome responses to the short-term effects of Amazonian deforestation. Molecular Ecology, 24(10), 2433–2448. http://doi.org/10.1111/mec.13172

Tin, H. S., Palaniveloo, K., Anilik, J., Vickneswaran, M., Tashiro, Y., Vairappan, C. S., & Sakai, K. (2018). Impact of Land-use Change on Vertical Soil Bacterial Communities in Sabah. Microbial Ecology, 75(2), 459–467. http://doi.org/10.1007/s00248-017-1043-6

Tripathi, B. M., Kim, M., Singh, D., Lee-Cruz, L., Lai-Hoe, A., Ainuddin, A. N., … Adams, J. M. (2012). Tropical Soil Bacterial Communities in Malaysia: PH Dominates in the Equatorial Tropics Too. Microbial Ecology, 64(2), 474–484. http://doi.org/10.1007/s00248-012-0028-8

Wood, S. A., Gilbert, J. A., Leff, J. W., Fierer, N., D’Angelo, H., Bateman, C., … McGuire, K. L. (2017). Consequences of tropical forest conversion to oil palm on soil bacterial community and network structure. Soil Biology and Biochemistry, 112, 258–268. http://doi.org/10.1016/j.soilbio.2017.05.019

